# Deep Mutational Scanning of Dynamic Interaction Networks in the SARS-CoV-2 Spike Protein Complexes: Allosteric Hotspots Control Functional Mimicry and Resilience to Mutational Escape

**DOI:** 10.1101/2021.06.15.448568

**Authors:** Gennady M. Verkhivker

## Abstract

We develop a computational approach for deep mutational scanning of residue interaction networks in the SARS-CoV-2 spike protein complexes to characterize mechanisms of functional mimicry and resilience to mutational escape by miniprotein inhibitors. Using a dynamic mutational profiling and sensitivity analysis of protein stability, binding interactions and global network parameters describing allosteric signaling, we identify regulatory hotspots in the SARS-CoV-2 S complexes with the ACE2 host receptor and ultra-potent miniproteins. The results revealed that global circulating variants are associated with allosteric control points that are dynamically coupled to structural stability hotspots. In this mechanism, variant-induced perturbations of flexible allosteric sites can result in global network changes and elicit specific protein responses. The binding affinity fingerprints and allosteric signatures of the SARS-CoV-2 complexes with miniproteins are determined by a dynamic cross-talk between regulatory control points and conformationally adaptable allosteric hotspots that collectively control structure-functional mimicry, signal transmission and resilience to mutational escape.

## Introduction

SARS-CoV-2 infection is transmitted when the viral spike (S) glycoprotein binds to the host cell receptor ACE2 leading to the entry of S protein into host cells and membrane fusion.^1,2^ Structural and biochemical studies established that the mechanism of virus infection may involve conformational transitions between distinct functional forms of the SARS-CoV-2 S protein between closed “down” and open “up” forms of the receptor-binding domain (RBD).^3-10^ Structural and functional studies of SARS-CoV-2 antibodies have examined molecular mechanisms underlying binding competition with the ACE2 host receptor, showing that antibodies often bind to the accessible regions of the SARS-CoV-2 S proteins where viruses can tolerate mutations and thereby escape immune challenge.^11-15^ Optimally designed combinations and synergistic cocktails of different antibody classes simultaneously targeting the conserved and more variable SARS-CoV-2 RBD epitopes can provide more efficient cross-neutralization effects and resilience against mutational escape.^16-19^ These studies also suggested that some ultra-potent antibodies may function by allosterically interfering with the host receptor binding and causing conformational changes in the S protein that can obstruct other epitopes and block virus infection without directly interfering with ACE2 recognition. Biochemical and functional studies using a protein engineering and deep mutational scanning have quantified binding mechanisms of SARS-CoV-2 interactions with the host receptor, showing that many mutations of the RBD residues can be well tolerated with respect to both folding and binding.^20,21^ Functional mapping of mutations in the SARS-CoV-2 S-RBD using deep mutational scanning showed that sites of antibody mutational escape are constrained by requirement of the protein stability and ACE2 binding, suggesting that escape-resistant antibody cocktails can limit the virus ability to acquire novel sites of immune escape in the RBD without compromising its binding to ACE2.^22-24^ Although functionally analogous to antibodies, nanobodies are much smaller and often more robust and thermostable. The recent investigation reported discovery of an ultra-potent synthetic nanobody Nb6 that neutralizes SARS-CoV-2 by stabilizing the fully inactive down S conformation preventing binding with ACE2 receptor.^25^Nonetheless, many potentially neutralizing nanobodies also recognize regions of RBD that are subject to escape variation, reducing their potential efficacy.^26-28^ Potent neutralizing nanobodies that resist convergent circulating variants of SARS-CoV-2 by targeting novel and conserved epitopes were recently reported.^26^ Cryo-EM structure determination of different classes of nanobodies revealed mechanisms of high-affinity and broadly neutralizing activity against the evolving virus by exploiting novel epitopes on the SARS-CoV-2 S protein that are not accessible to antibodies.^26^

The miniaturization of high-affinity protein binders for the SARS-CoV-2 S protein focused on design of protein mimics with enhanced stability and smaller sizes to take advantage over antibodies for direct delivery into the respiratory system. Protein engineering approaches combined with functional studies generated smaller neutralizing proteins characterized by low nanomolar to femtomolar binding affinities with SARS-CoV-2 S proteins, including nanobody Nb6,^25^ monoclonal antibody 2B04,^29^ protein decoy mimics,^30^ and mini-protein inhibitors.^31^ De novo design engineering of decoy proteins that bind to the SARS-CoV-2 S protein by replicating the ACE2 interface produced low nanomolar affinity monovalent decoy CTC-445.2 with high specificity to the S-RBD region.^30^ Cryo-EM structure determination confirmed that CTC-445.2 can simultaneously bind to all three RBDs of the SARS-CoV-2 S protein showing structural mimicry with the ACE2 binding and thereby suggesting that mutational escape against decoy binding would be unlikely. Another illuminating study reported de novo design of high affinity mini-protein inhibitors using computer-generated scaffolds to optimize binding, folding, and stability of the SARS-CoV-2 S protein.^31^ Cryo-EM structures and biochemical studies of hyper-stable minibinders in complexes with the SARS-CoV-2 S ectodomain trimer confirmed computational predictions and revealed picomolar binding affinity with neutralization of the SARS-CoV-2 that is superior to the most potent monoclonal antibodies.^31^ The high and specific binding affinity of the optimized de novo protein decoys also lead to an effective and specific in vitro neutralization of SARS-CoV-2 viral infection. Despite general resistance of mini-proteins to viral mutations in the RBD-binding interface, the experimental studies demonstrated that mutational escape could still be possible when multiple mutations that have a greater differential binding effect on are combined.

The important questions that could provide further impetus towards rational design of protein binders for the SARS-CoV-2 S target are: (a) characterization of allosteric molecular signatures of sites targeted by global viral variants; and (b) understanding how structural mimicry and protein miniaturization modulate allosteric communication footprint and whether these changes affect resilience towards mutational escape. In this study, we propose a computational approach for deep mutational profiling of the SARS-CoV-2 S protein binding and allosteric interaction networks to characterize mechanisms of structural mimicry and mutational escape in the SARS-CoV-2 complexes. Our approach is built upon previous computational simulation studies of the SARS-CoV-2 spike proteins and complexes^32-34^ and particularly a series of our investigations suggesting that SARS-CoV-2 S proteins are functionally adaptable and allosterically regulated machines that exploit the intrinsic plasticity of functional regions and versatility of allosteric hotspots to modulate complex phenotypic responses to the host receptor and antibodies.^35-39^ Through mutational scanning of binding energetics and perturbation-based approach for profiling of dynamic interaction networks, we identify binding affinity fingerprints and allosteric signatures of the SARS-CoV-2 complexes with the ACE2 host receptor and a group of miniprotein binders. We show that while stable regulatory centers enable structural mimicry in the SARS-CoV-2 complexes, mutational variants and escape mechanisms may be associated with conformationally adaptable allosteric hotspots that are dynamically coupled to stability centers, thereby allowing for local perturbations in these flexible positions to induce functional allosteric changes in the residue interaction networks and elicit fine-tuning of the protein response and activity.

## Results and Discussion

### Dynamic Signatures of the SARS-CoV-2 S Complexes with Miniprotein Binders

We performed all-atom MD simulations of the SARS-CoV-2 structures with ACE2, CTC-4452 (Figure 1A,D), LCB1 (Figure 1B,E) and LCB3 proteins (Figure 1C,F) and the equilibrium conformational ensembles were employed for a distance fluctuations coupling analysis of the dynamic residue correlations to probe stability and allosteric communication preferences of the SARS-CoV-2 S residues. The residue-based distance fluctuation stability index measures the energy cost of the dynamic residue deformations and can serve as a robust metric for an assessment of both structural stability and allosteric propensities of protein residues. In this model, dynamically correlated residues whose effective distances fluctuate moderately are expected to communicate with the higher efficiency than the residues that experience large fluctuations. Notably, densely interconnected residues with high value of these indexes often serve as structurally stable centers and regulators of allosteric signals. The results displayed a strong similarity between the dynamic fluctuation profiles of the SARS-CoV-2 S-RBD complexes with ACE2 and miniprotein binders (Figure 2A-C). The dominant peaks of the profiles corresponded to structural stability centers F400, I402, Y505, Y508, F490, Y489, and A475 and are preserved in all complexes.

**Figure 1.**
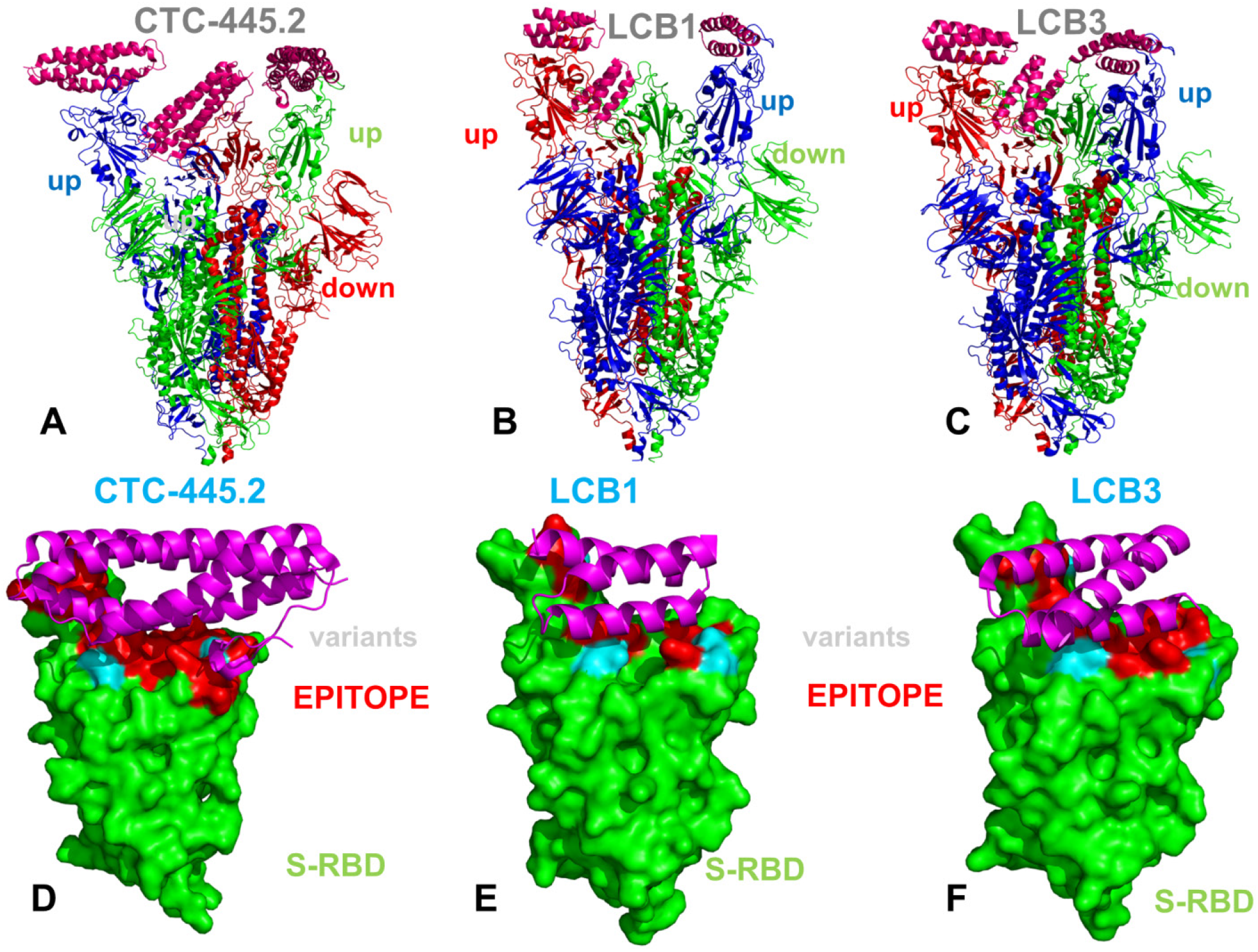
Cryo-EM structures of the SARS-CoV-2 S trimer structures used in this study. The structures of SARS-CoV-2 S trimer in the complex with the ACE2 protein decoy CTC-445.2, pdb id 7KL9 (A), in the complex with the designed miniprotein binder LCB1, pdb id 7JZL (B) complex with the miniprotein LCB3, pdb id 7JZN (C). The structure is in ribbons with protomers A,B,C are colored in green, red and blue respectively. The S trimer complexes with CTC-445.2, LCB1 and LCB3 have two protomers in the up position and one protomer in the down-closed position. The respective protomer up/down orientation is annotated. The binding epitopes of the S-RBD bound structures are shown for CTC-445.2, LCB1 and LCB3 in panels (D-F) respectively. S-RBD is in green surface and binding epitopes are in red. The bound miniproteins are shown in magenta-colored ribbons and annotated. The epitope binding residues are shown in red surface. The positions of functional sites K417, L452, E484, and N501 subjected to mutational circulating variants are shown in cyan.

**Figure 2.**
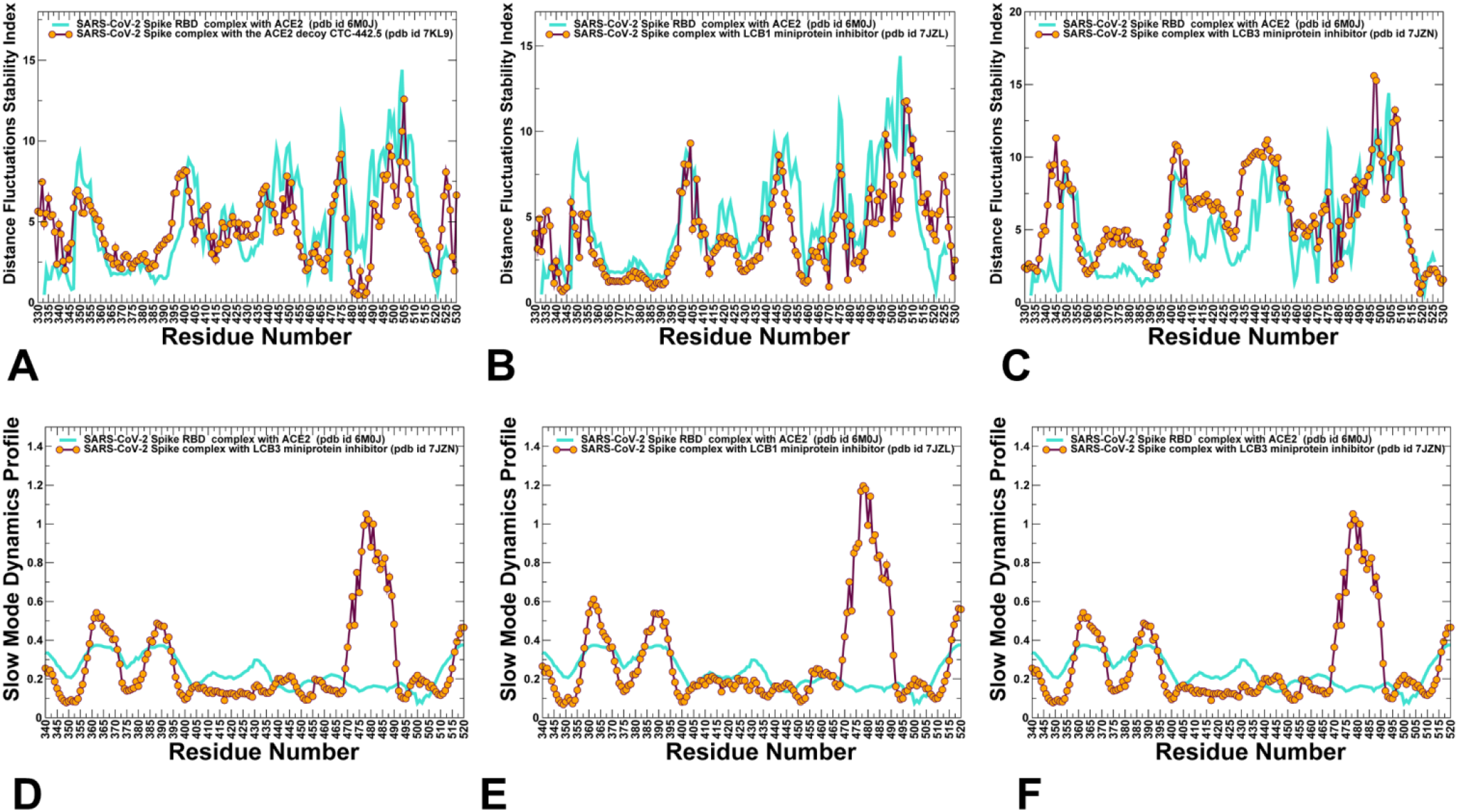
The distance fluctuation stability index for the SARS-CoV-2 S complexes with CTC-445.2 (A), LCB1 miniprotein (B), and miniprotein LCB3 (C). The profiles are shown for S-RBD residues in maroon-colored lines with orange-filled circles. The respective profile for the S-RBD complex with ACE2 is shown in turquoise colored lines. Collective dynamics profiles of the SARS-CoV-2 S complexes are shown for CTC-445.2, LCB1 and LCB3 protein binders in panels (D-F) respectively. The profiles represent the mean square displacements in functional motions averaged over the three lowest frequency modes. The slow mode profiles are shown in maroon-colored lines with individual data points highlighted in orange circles. In each panel, the distributions are shown together with the slow mode profile for the S-RBD complex with ACE2 (in turquoise color lines).

While the shapes of the dynamic profiles are similar in the SARS-CoV-2 S complexes with ACE2 and miniproteins, there are important local differences that reflect the binding-induced flexibility changes (Figure 2A-C). The dynamic fluctuation stability profile featured several moderate local peaks near K417/I418 and L452/Y453 positions and a considerably stronger peak corresponding to residues N501/Y505 for the RBD-ACE2 complex. The conservation of communication clusters near these functional positions is mainly determined by stability of hydrophobic residues I418, L453 and L455 that are dynamically coupled to more flexible K417 and L452 sites. Notably, the stability peak near N501 residue induced by binding in the RBD-ACE2 complex may be partly weakened in the complexes with miniproteins (Figure 2A-C). At the same time, E484 belongs to a flexible RBD region with small values of distance fluctuation couplings index, implying that thermal fluctuations in this position could be only weakly coupled to perturbations of other protein residues. Importantly, the results suggested that structural mimicry between binding interfaces formed by the ACE2 host receptor and miniproteins can result in respective considerable similarities of the dynamic correlation inter-residue couplings in the functional regions. The distance fluctuation profiles suggested that binding of miniproteins can also induce similar changes in the distribution of rigid and flexible S-RBD regions as the host receptor ACE2 (Figure 2A-C).

Structural maps of conformational mobility profiles of the SARS-CoV-2 S trimer complexes further illustrated that the ACE2 protein decoy and miniprotein binders result in similar dynamic characteristics and patterns of stability and flexibility (Supporting Information, Figure S1). As expected, the S2 subunit regions (residues 910-1163) showed considerable structural rigidity, while S-RBD regions are more dynamic, especially in the protomers assuming open conformation. Structural and dynamic similarities are also evident from the covariance matrixes of residue fluctuations suggesting conservation of correlated motions (Supporting Information, Figure S1). These maps are consistent with the distance fluctuation correlation profiles, indicating that structural and more flexible residues in the S-RBD binding interface may be dynamically coupled in the SARS-CoV-2 S protein complexes with miniproteins.

We also characterized collective motions for the SARS-CoV-2 complexes using principal component analysis (PCA) of the MD trajectories (Figure 2D-F). The slow functional modes of dynamic residue fluctuations are determined primarily by the protein fold topology and play an important role in synchronizing collective motions and controlling allosteric conformational changes. Allosteric protein responses can be induced when mutations or interacting protein partners can exploit the energetically favorable movements along the slow modes.^40,41^ The overall shape of the essential profiles was generally preserved between SARS-CoV-2 S complexes with ACE2 and miniproteins (Figure 2D-F). We noticed that functional positions of circulating variants K417 and L452 belong to regions associated with local minima, pointing to potential hinge clusters involved in regulation of collective motions. A striking difference in the slow mode profiles lies in the flexible region centered on the E484/F486 RBD positions. While in the complex with ACE2 the mobility of this intrinsically flexible RBD region is considerably reduced, these key functional sites corresponded to strong maxima of the essential mobility profiles in the S complexes with miniproteins (Figure 2D-F). As a result, this flexible region can experience large scale functional movements and contribute to allosteric conformational changes induced by binding of miniprotein inhibitors. A much smaller peak of the distribution is aligned with N501 position, indicating some level of functional motions along slow modes. Hence, sites E484 and N501 targeted by circulating variants belong to the functional regions undergoing concerted motions along pre-existing slow modes of the S protein. Interestingly, binding of miniproteins could preserve the dynamic signature of the unbound spike protein in this region, which can ensure resilience of these inhibitors to mutational escape. To further explore similarities in structural and dynamic signatures of the SARA-CoV-2 S trimer complexes, we used equilibrium trajectories to compute the ensemble-averaged solvent accessible surface area (SASA) residue profiles for in the SARS-CoV-2 RBD complexes flexibility (Supporting Information, Figure S2). Consistent with a high conservation and structural stability of the S2 subunit, the distributions showed that the inter-protomer interface residues are generally buried in the core of the S2 subunit. The S-RBD core residues and binding interface S residues showed relatively moderate SASA values. Of particular interest was analysis of patterns exhibited by the functional sites targeted by circulating variants. In the S trimer complex with CTC-445.2 decoy, K417 and E484 are fairly exposed and flexible, while L452 is partially buried, and N501 is almost entirely buried at the binding interface (Supporting Information, Figure S2A). In the complexes with miniproteins LCB1 and LCB3, only E484 is exposed to solvent, while other functional sites are partly buried but could retain some degree of dynamic plasticity (Supporting Information, Figure S2B,C).

### Deep Mutational Scanning Identifies Structural Stability and Binding Affinity Hotspots in the SARS-CoV-2 Complexes: Conserved Hotspots Enable Functional Mimicry for Miniprotein Inhibitors

Using the ensemble-based mutational scanning of protein stability and binding affinity, we constructed mutational heatmaps and compared the binding energy hotspots in the SARS-CoV-2 complexes with ACE2 (Figure 3A), CTC-445.2 decoy (Figure 3B), LCB1 and LCB3 mini-proteins (Figure 3C,D). The results of mutational scanning produced G446, Y449, Y453, L455, F456, F486, Q493, Y489, and Y505 as key binding energy hotspots for ACE2 binding (Figure 3A). These data are consistent with deep mutagenesis scanning studies that have detailed SARS-CoV-2 interactions with the ACE2 host receptor and key energetic hotspots of binding and stability.^20-22^ Mutational sensitivity analysis of SARS-CoV-2 RBD binding with CTC-445.2 decoy revealed a strong similarity in the distribution of binding energy hotspots, showing that structural and dynamic mimicry resulted in the preservation of ACE2-binding patterns, also supporting the experimental findings of CTC-445.2 resilience to viral mutations (Figure 3B). The key binding hotspots Y453, L455, F456, and Y489 are shared between ACE2 and decoy binding. It is worth noting that mutational scanning confirmed small changes in the binding affinity caused by modifications in functional positions subjected to circulating variants. Moreover, binding affinity of the CTC-445.2 protein decoy appeared to be insensitive to variations in K417, L452 and E484 positions. The important finding of deep mutational scanning is a consistent pattern of conserved energy hotspots is preserved in the complexes with miniprotein inhibitors.

**Figure 3.**
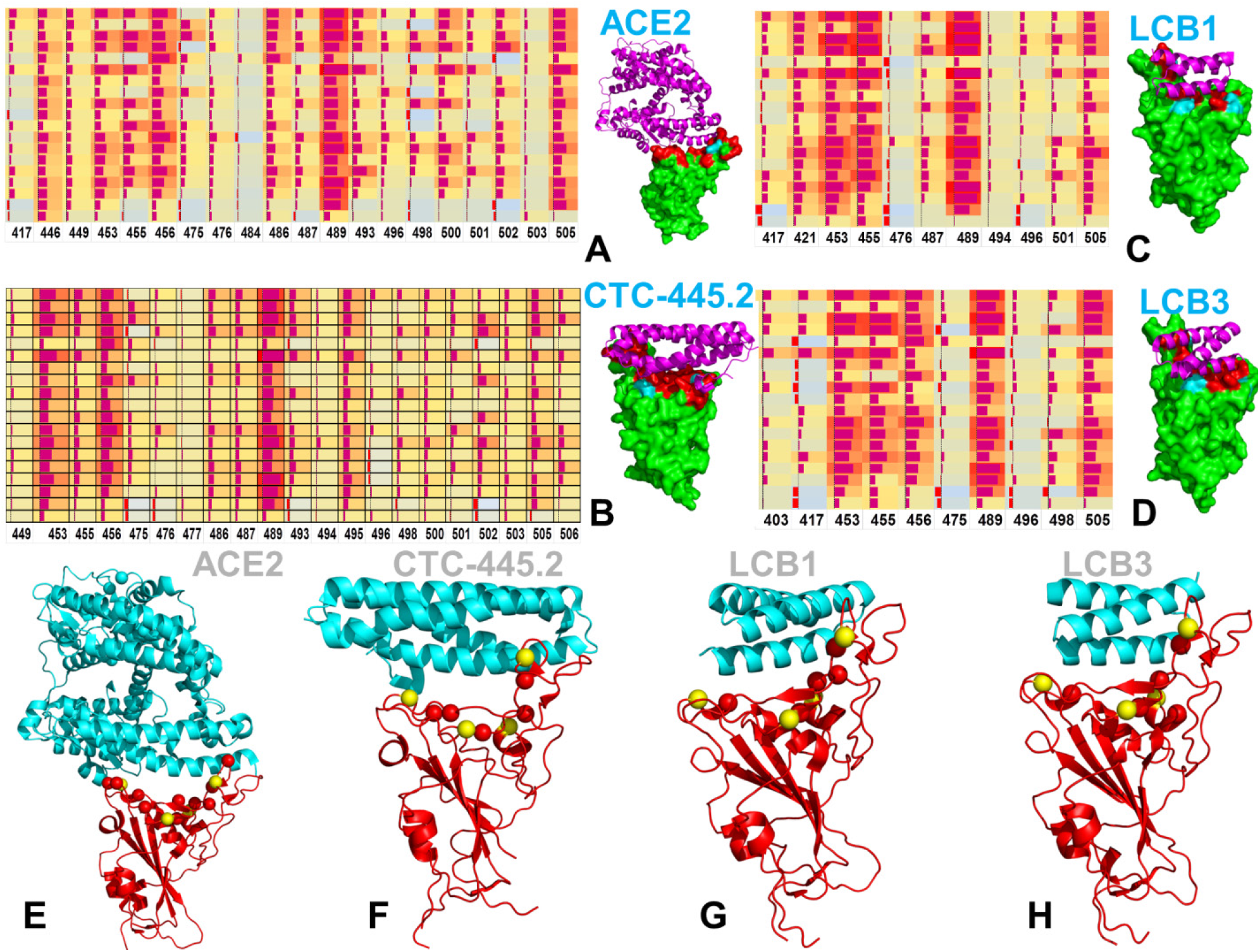
The mutational scanning heatmaps for the SARS-CoV-2 S complexes with ACE2, pdb id 6M0J (A), ACE2 protein decoy CTC-445.2, pdb id 7KL9 (B), and miniproteins LCB1, pdb id 7JZL (C) and LCB3, pdb id 7JZN (D). The heatmaps show the computed binding free energy changes for 19 single mutations on the binding epitope sites. The squares on the heatmap are colored using a 3-colored scale - from light blue to red, with red indicating the largest destabilization effect. The data bars correspond to the binding free energy changes, with positive destabilizing value shown by bars towards the right end of the cell and negative stabilizing values s bars oriented towards left end. The length of the data bar is proportional to the value in the square cell. S-RBD is shown in green surface. The binding epitope regions are shown in red. The positions of K417, L452, E484 and N501 are shown in light cyan color and annotated. Structural mapping of the binding energy hotspots is shown for the SARS-CoV-2 S complexes with ACE2 (D), CTC-445.2 (E), LCB1 (F) and LCB3 (G). The binding energy hotspots are in red spheres and correspond to residues with high mutational sensitivity. The functional sites K417, L452, E484, and N501 are in yellow spheres.

Mutational heatmaps with the miniprotein binders LCB1 and LCB3 revealed that the binding hotspots Y421, Y453, L455, F456, Y489 and Y505 are shared with ACE2 (Figure 3C,D). Strikingly, despite a vastly different size of the binding interface as compared to ACE2, the bulk of the binding free energy can be provided by a small number of critical hotspots responsible for both protein stability and binding affinity. The mutational scanning analysis also highlighted the energetic efficiency of small binding interfaces formed by miniproteins LCB1/LCB3 where remarkably essentially all residues from the binding epitope can serve as strong binding hotspots (Figure 3C,D). Coupled with a relatively small size and arguably small entropic losses upon binding for miniproteins, this can rationalize the experimentally observed ultra-potency of LCB1 and LCB3 inhibitors that surpasses binding affinities of the most potent monoclonal antibodies.^31^ According to mutational heatmaps, mutations of N501 produced minor destabilization changes on the binding affinity with the CTYC-445.2 decoy, while some mutations of this functional site could lead to the improved binding affinity with ACE2 (Figure 3). In addition, small differences were observed in positions G446, Y449, F486 and Y505 that are less significant in the complex with CTC-445.2 while the contribution of Y495 residue becomes appreciably stronger than in the complex with ACE2 (Figure 3A,B). Mutational cartography also demonstrated only small binding energy changes caused by modifications in sites of circulating variants (Figure 3B-D). Combined with the functional dynamics analysis, these results suggested that the miniprotein binders LCB1 and LCB3 can be resilient to mutational escape as binding energetics and collective motions appeared to be relatively tolerant to perturbations in these functional sites.

The computed strong correlation between S-RBD binding free energy changes with ACE2 and CTC-445.2 (Pearson correlation coefficient R = 0.85) demonstrated that the decoy can leverage not only key binding hotspots but also remarkably replicate the entire nature binding interface (Figure 4A). The results are in excellent agreement with the experimental deep mutational scanning.^30^ Indeed, site saturation mutagenesis (SSM) experiments compared the effect of single–amino acid substitution on binding to ACE2 or CTC-445.2 decoy, yielding a similar correlation between binding to the ACE2 host receptor and CTC-445.2 decoy (Pearson correlation coefficient R = 0.92).^30^ The results and obtained agreement with the experiment supported the notion that CTC-445.2 protein decoy can maintain resiliency to escaping mutations (Figure 4A).

**Figure 4.**
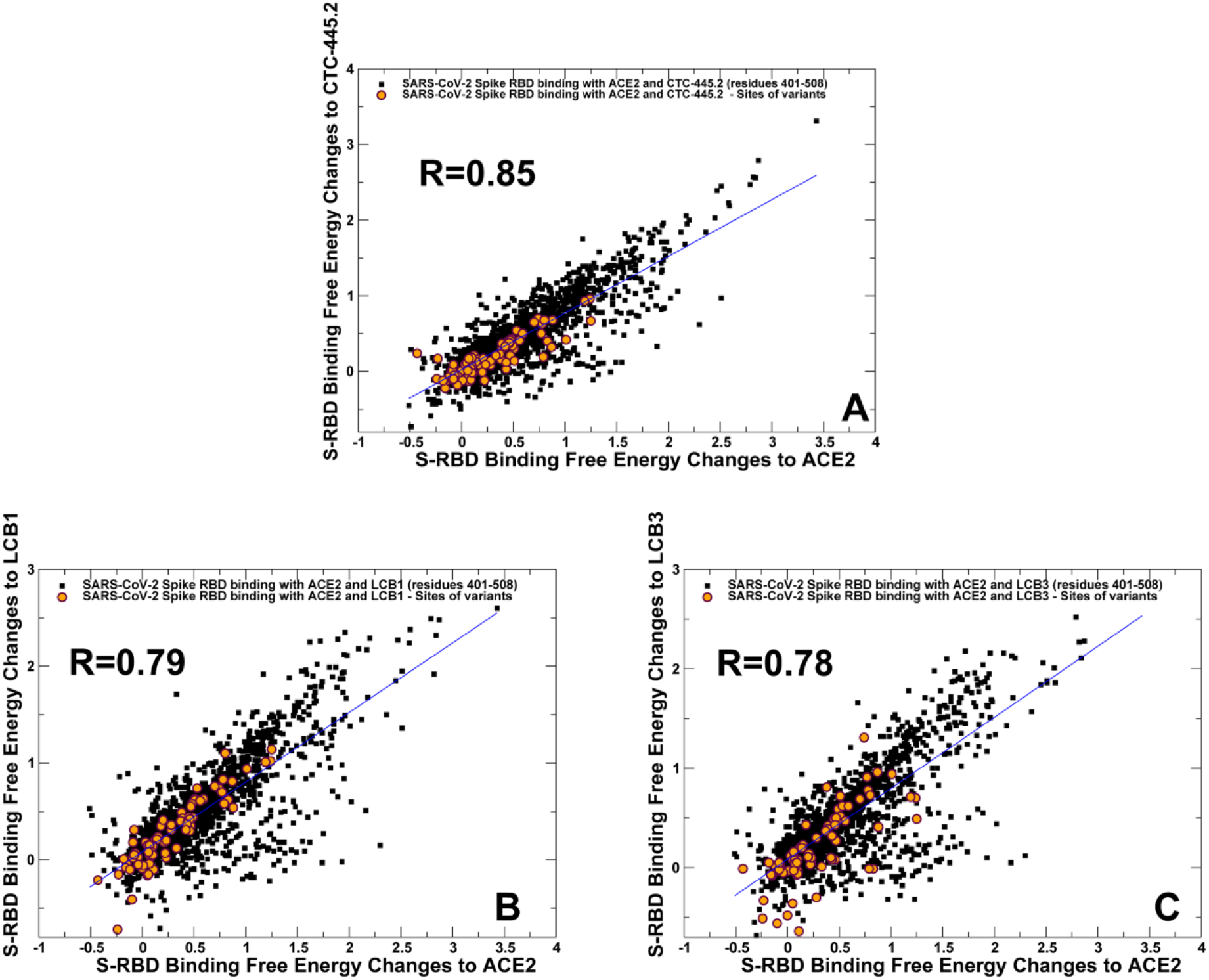
The 2D scatter plots of binding free energy changes for the S-RBD residues in complexes with ACE2 and miniproteins. (A) The scatter plot of binding free energy changes from deep mutational scanning of the S-RBD residues with ACE2 and CTC-445.2 (Pearson correlation coefficient R = 0.85). (B) The scatter plot of binding free energy changes from deep mutational scanning of the S-RBD residues with ACE2 and LCB1 (Pearson correlation coefficient R = 0.79). (C) The scatter plot of binding free energy changes from deep mutational scanning of the S-RBD residues with ACE2 and LCB3 (Pearson correlation coefficient R = 0.78). The data points are shown as black-filled squares and scatter points corresponding to the binding free energy changes in sites K417, L452, E484 and N501 are shown as orange-filled circles. A consistent high correlation between the effects of mutations in the RBD residues on binding highlights the mutational resilience of the miniprotein analogues.

Of particular interest were the binding free energy changes associated with modifications at sites of circulating variants K417, L452, E484, and N501 (Figure 4). A similarly strong correlation of the binding free energy changes in these positions was seen for the SARS-CoV-2 S complexes with ACE2 and CTC-445.2 decoy (Figure 4A). Notably, the mutation-induced energetic changes in these positions were moderate, with most of these differences smaller than 1.0 kcal/mol (Figure 4A). Hence, structural and dynamic replication of the protein stability and binding interactions by the CTC-445.2 decoy could potentially curtail the escaping “opportunities” for the virus as mutations that would impede or reduce binding with the ACE2 decoy would also adversely affect protein stability and binding with the host receptor.

These findings showed structural and energetic mimicry of the protein decoy is directly linked with the resilience to mutational escape allowing for effective and specific neutralization of SARS-CoV-2 viral infection. Nonetheless, despite and overall robust resiliency to viral mutations in the RBD-binding interface, some mutations in positions G447, N487 and Y495 were more sensitive to CTC-445.2 binding and had smaller effect on binding with ACE2. At the same time, mutations of N487 residue yielded moderate destabilization changes and could arguably induce only minor loss of binding affinity (Figure 4A). The experimental deep mutational profiling of the S-RBD residues indicated that viral escape mutations are consistently those that have large effects on antibody or protein binding escape but elicit only minor impact on ACE2 binding and S-RBD folding stability.^23,24^ As residues G447 and Y495 corresponded to structurally stable centers, mutations in these positions are likely to negative impair the RBD stability and therefore unlikely to emerge as viable escape variants under physiological conditions. These results suggested that structural mimicry of the protein decoy can ensure the intrinsic resiliency to mutational escape as mutations imposing little cost on the RBD folding or ACE2 binding affinity would be readily tolerated in the complex the protein decoy.

A comparison of mutational effects on the binding free energies with the LCB1 (Figure 4B) and LCB3 miniproteins (Figure 4C) also yielded a similar pronounced correlation with Pearson’s coefficient R=0.796 and R=0.784 respectively. The changes associated with mutations in K417, L452, E484, and N501 positions showed a somewhat larger spread of binding free energies, particularly for modifications predicted to improve binding affinity that are larger for miniproteins (Figure 4B,C).

### Network-Based Mutational Profiling of Allosteric Signaling: Dynamic Crosstalk of Structural Stability Centers and Functionally Adaptable Allosteric Hotspots Controls Resilience to Mutational Escape

We adapted a dynamic model of residue interaction networks for deep mutational scanning of allosteric interactions by computing mutation-induced changes in the topological network parameters that characterize allosteric signaling. Through mutational perturbations of the S-RBD positions we reevaluate dynamic inter-residue couplings and compute changes in the average short path length (ASPL) that are averaged over all possible substitutions in a given residue node. These global network characteristics are used to characterize the average mutational effect of each node on changes in the network efficiency and allosteric communications. By identifying residues where mutations trigger a significant change of the global topological parameter, this network-based mutational profiling identifies allosteric hotspot positions that are important for signal transmission in the SARS-CoV-2 complexes. The distribution of mutation-induced ASPL changes in the complexes with the ACE2 (Figure 5A,D) and CTC-445.2 (Figure 5B,E) pointed to distinct residue clusters characterized by a significant increase of the ASPL parameter. Mutations of these positions on average lead to the increased path length and therefore refer to sites that may be important for efficiency of signal transmission in the complexes. The first cluster is formed by the S-RBD core residues (S383, K386, and N388), the second cluster (L452, Y453, L455, F456) is anchored around structural stability center Y453, and cluster (Y495, W498, Y489, N501, Y505) included stability hotspots Y489 and Y505 (Figure 5A). Hence, mutations of the binding energy hotspots Y453, L455, Y489 and Y505 could also result in a reduction of network efficiency, confirming the essential role of these regulatory centers in propagation of allosteric signals in the complex.

**Figure 5.**
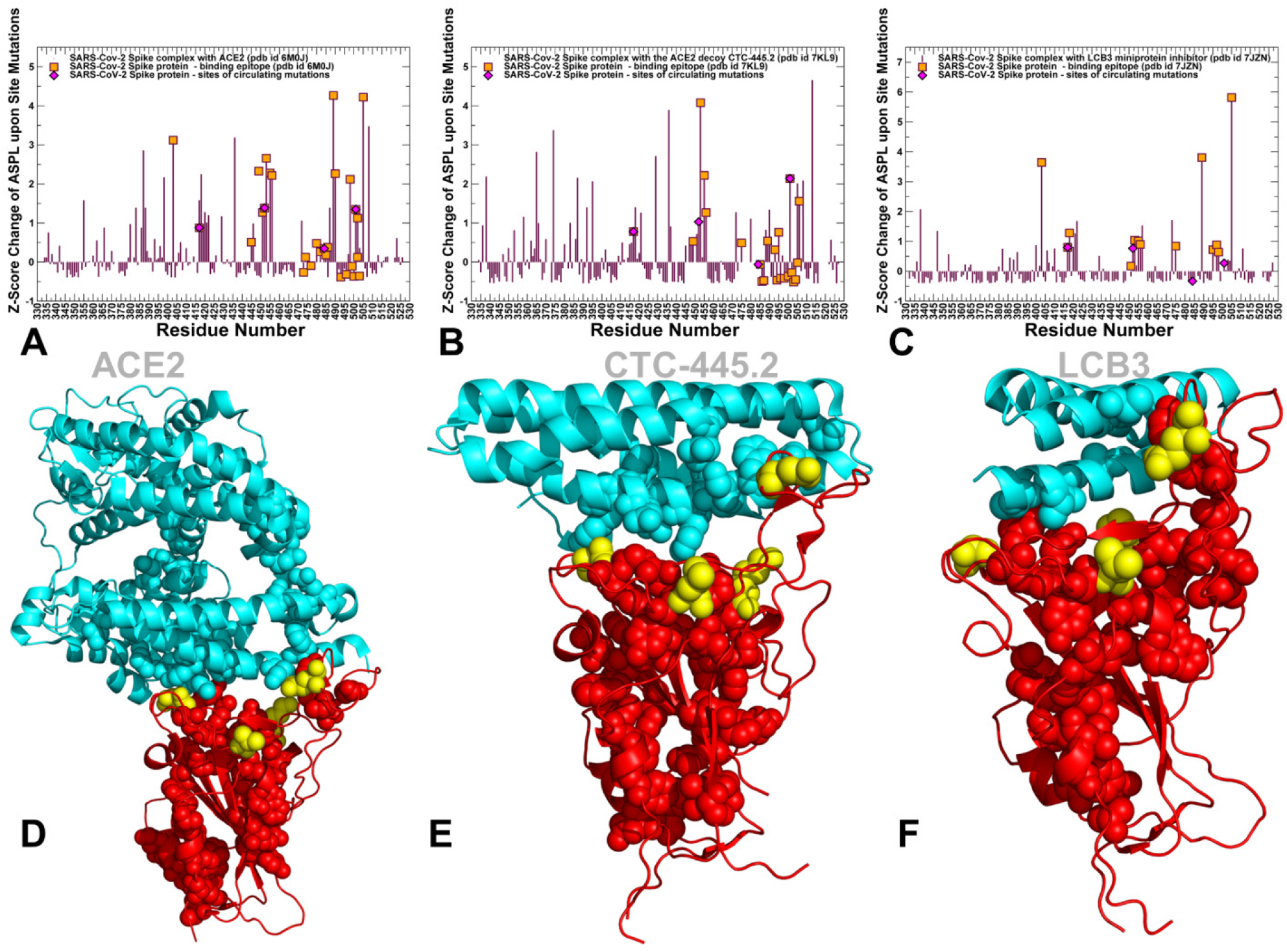
Mutation-averaged changes in the topological network parameters for the SARS-CoV-2 S-RBD complexes. The residue-based Z-score profile measuring mutational change in the ASPL parameter for the S-RBD complex with ACE2 (A), complex with CTC-445.2 (B), and LCB3 miniprotein binder (C). The global ASPL parameter is recalculated upon mutational scanning of single nodes and the average Z-score change is estimated for each residue node. The profiles are shown as maroon-colored bars. The positions of binding epitope residues are shown in orange-filled squares, and the positions of functional sites K417, L452, E484 and N501 are shown as magenta-filled diamonds. Structural maps of the probably network communication routes are shown for the S-RBD complex with ACE2 (D), complex with CTC-445.2 (E), and LCB3 miniprotein binder (F). The paths are formed by residues with the high edge betweenness values which is an edge centrality parameter related to the number of shortest paths linking any pair of residues in the network and passing through a given edge. The S-RBD is shown in red ribbons in panels (D-E). The high edge betweenness residues are in red-colored spheres. The functional sites K417, L452, E484, and N501 are in yellow spheres. The binding proteins ACE2, CTC-445.2 and LCB3 are shown in cyan colored ribbons and annotated.

Strikingly, the sites of circulating variants K417, L452 and N501 were prominently featured among these clusters in the S-RBD complex with ACE2, although the respective ASPL changes were smaller for these dynamic positions than for the structural stability centers (Figure 5A). We also observed that Y453 and Y508 residues corresponded to the major regulatory centers that control propagation of allosteric communication between the S-RBD and CTC-445.2 decoy (Figure 5B). Another two major peaks are F429, I418, F490, Y495, N501 and Y508 residues. In addition, a conserved group in the RBD core (F374, S383, L387) also featured high peaks, as mutations in these sites could impair both protein stability and allosteric couplings.

Notably, N501 emerged as one of the major allosteric hotspots of the S-RBD complex with the CTC-445.2 decoy (Figure 5B). Mutation-induced allosteric changes upon modifications of K417 and L452 positions were also appreciable, while mutations in E484 showed only a minor effect on network changes. These results suggested that through dynamic coupling with neighboring hotspots I418 and Y453, modifications of K417 and L452 sites could modulate allosteric interactions and communications between S-RBD and CTC-445.2 decoy (Figure 5B). Hence, despite structural mimicry between ACE2 and the protein decoy, mutational escape effects to CTC-445.2 binding may still arise due to some modifications in the N501 and L452 positions that could affect efficiency of allosteric signaling in the complex.

Structural mapping of high edge betweenness links highlighted the probable routes of allosteric communication, showing that sites of circulating variants could bridge communication routes and mediate their access to the binding interface (Figure 5D,E). The residues that featured significant Z-score changes in ASPL upon mutations were also mapped onto SARS-CoV-2 S-RBD structures (Supporting Information, Figure S3). These maps illustrated conservation of the important allosteric sites in the S-RBD core and several allosteric communication centers in the RBD binding interface (Supporting Information, Figure S3).

Importantly, functional sites K417, L452 and N501 are dynamically coupled with major structural stability hotspots, yielding distinct allosteric hotspot clusters that together can control signal transmission (Figure 5, Supporting Information, Figure S3). As sites of circulating variants are less conserved and more flexible, their dynamic coupling to the regulatory centers of structural stability could allow for global changes in the residue interaction networks and functional modulation of signal transmission. We argue that mutations accompanied by moderate perturbations in these positions could cascade to major functional sites and allosterically modulate functional dynamics. This may underlie a plausible mechanism by which minimal perturbations in mutational escape positions may induce global protein response by allosterically altering the interaction networks near the functionally critical sites.

In the complex with LCB1 and LCB3 miniproteins, the largest changes upon mutations in the network parameter distribution occur at positions Y505, Y489, F400, I418, N422, F497, Y453 and I418 (Figure 5C,F). These residues with the exception of Y489 and Y453 do not correspond to the binding free energy hotspots, but may be important mediating sites of the allosteric interaction network. The ASPL changes caused by mutations in K417 and L452 are moderate and for N501 and E484 positions are very small (Figure 5C,F). Structural analysis of communication routes highlighted a very similar alignment of major paths in S-RBD complexes with ACE2 and miniprotein. The circulating variant sites are dynamically coupled to major communication routes, with L452 and K417 residues immediately adjacent to key allosteric hotspots Y453 and I418. Our analysis suggested that mutations in these positions would likely have only minor effect on allosteric signaling, thus ensuring preservation of the inhibitor resilience against mutational escape caused by circulating variants.

To conclude, through comparison of the mutational scanning heatmaps and mutation-induced changes in the global network parameters, we showed that binding affinity fingerprints and allosteric signatures of the SARS-CoV-2 complexes with ACE2 and high affinity miniprotein mimics may be determined by communicating functional hotspots to preserve stability, ensure adaptation to mutational escape and provide for efficient allosteric signaling. The results suggested a mechanism in which resilience to mutational escape may be controlled by flexible allosteric hotspots that are dynamically coupled to critical for binding and stability positions, thereby allowing for cascading changes in the residue interaction network and effective fine-tuning of the protein response. Through this mechanism, SARS-CoV-2 spike protein may generate escape mutants to modulate binding and allosteric communications with protein partners.

## Methods

### Structure Preparation and Analysis

The cryo-EM structures of SARS-CoV-2 S proteins used in this study included the S-RBD complex with ACE2 (pdb id 6M0J), and SARS-CoV-2 S trimer complexes with the ACE2 protein decoy CTC-445.2 (pdb id 7KL9), S trimer complex with the designed miniprotein binder LCB1 pdb id 7JZL) and miniprotein LCB3 (pdb id 7JZN) (Figure 1). All structures were obtained from the Protein Data Bank.^42,43^ Hydrogen atoms and missing residues were initially added and assigned according to the WHATIF program web interface.^44,45^ The structures were further pre-processed through the Protein Preparation Wizard (Schrödinger, LLC, New York, NY) and included the check of bond order, assignment and adjustment of ionization states, formation of disulphide bonds, removal of crystallographic water molecules and co-factors, capping of the termini, assignment of partial charges, and addition of possible missing atoms and side chains that were not assigned in the initial processing with the WHATIF program. The missing loops in the cryo-EM structures were also reconstructed using template-based loop prediction approaches ModLoop^46,47^ and ArchPRED.^48^ The side chain rotamers were refined and optimized by SCWRL4 tool.^49^ The protein structures were subsequently optimized using atomic-level energy minimization with a composite physics and knowledge-based force fields using 3Drefine method.^50^

The shielding of the receptor binding sites by glycans is an important feature of viral glycoproteins, and glycosylation on SARS-CoV-2 proteins can camouflage immunogenic protein epitopes.^51,52^ While all-atom MD simulations with the explicit inclusion of the complete glycosylation shield provide a rigorous and the most detailed description of the conformational landscape for the SARS-CoV-2 S proteins, such simulations remain to be computationally highly challenging due to the size of a complete SARS-CoV-2 S system. The structure of glycans at important glycosites (N122, N165, N234, N282, N331, N343, N603, N616, N657, N709, N717, N801, N1074, N1098, N1134, N1158) of the closed and open states of SARS-CoV-2 S protein was previously determined in the cryo-EM structures of the SARS-CoV-2 spike S trimer in the closed state (pdb id 6VXX) and open state (pdb id 6VYB). These glycans were incorporated in simulations of the SARS-CoV-2 S protein complexes in addition to the experimentally resolved glycan residues present in the cryo-EM structures of studied SARS-CoV-2 S proteins.

All crystallographic water molecules and other heteroatoms including zinc were initially removed in simulations of the S-RBD complex with ACE2. The zinc-binding site is located near the bottom of the cleft of subdomain I of ACE2 and more than 20 Å away from the intermolecular binding interface with SARS-CoV-2 RBD proteins. Furthermore, previous studies have established that zinc does not stabilize ACE2 protein structure since the native and apo-enzymes are equally susceptible to heat denaturation.^53^ In addition, it was shown that metal replacements mainly affect catalytic activity while a negligible change on ACE2 binding and dissociation constant can be observed.^53,54^

### MD Simulations

All-atom 1µs MD simulations have been performed for all studied protein structures. The structures of the SARS-CoV-2 S trimers were simulated in a box size of 85 Å × 85 Å × 85 Å with buffering distance of 12 Å. Assuming normal charge states of ionizable groups corresponding to pH = 7, sodium (Na+) and chloride (Cl-) counter-ions were added to achieve charge neutrality and a salt concentration of 0.15 M NaCl was maintained. All Na^+^ and Cl^-^ ions were placed at least 8 Å away from any protein atoms and from each other. All-atom MD simulations were performed for an N, P, T ensemble in explicit solvent using NAMD 2.13 package^55^ with CHARMM36 force field.^56^ Long-range non-bonded van der Waals interactions were computed using an atom-based cutoff of 12 Å with switching van der Waals potential beginning at 10 Å. Long-range electrostatic interactions were calculated using the particle mesh Ewald method^57^ with a real space cut-off of 1.0 nm and a fourth order (cubic) interpolation. Simulations were run using a leap-frog integrator with a 2 fs integration time step. All atoms of the complex were first restrained at their crystal structure positions with a force constant of 10 Kcal mol^-1^ Å^-2^. Equilibration was done in steps by gradually increasing the system temperature in steps of 20K starting from 10K until 310 K and at each step 1ns equilibration was done keeping a restraint of 10 Kcal mol-1 Å-2 on the protein C_α_ atoms. After the restrains on the protein atoms were removed, the system was equilibrated for additional 10 ns. An NPT production simulation was run on the equilibrated structures for 500 ns keeping the temperature at 310 K and constant pressure (1 atm). In simulations, the Nose–Hoover thermostat^58^ and isotropic Martyna–Tobias–Klein barostat^59^ were used to maintain the temperature at 310 K and pressure at 1 atm respectively.

### Functional Dynamics and Collective Motions

We performed principal component analysis (PCA) of reconstructed trajectories derived from CABS-CG simulations using the CARMA package^60^ and also determined the essential slow mode profiles using elastic network models (ENM) analysis.^61-63^ Two elastic network models: Gaussian network model (GNM) and Anisotropic network model (ANM) approaches were used to compute the amplitudes of isotropic thermal motions and directionality of anisotropic motions. The functional dynamics analysis was conducted using the GNM in which protein structure is reduced to a network of *N* residue nodes identified by *C*_*α*_ atoms and the fluctuations of each node are assumed to be isotropic and Gaussian. Conformational mobility profiles in the essential space of low frequency modes were obtained using ANM server^63^ and DynOmics server.^64^

### Local Structural Parameters: Relative Solvent Accessibility

The relative solvent accessibility parameter (RSA) was computed that is defined as the ratio of the absolute solvent accessible surface area (SASA) of that residue observed in a given structure and the maximum attainable value of the solvent-exposed surface area for this residue.^65^ According to this model, residues are considered to be solvent exposed if the ratio value exceeds 50% and to be buried if the ratio is less than 20%. Analytical SASA is estimated computationally using analytical equations and their first and second derivatives and was computed using web server GetArea.^65^

### Distance Fluctuations Coupling Analysis

We employed distance fluctuation analysis of the simulation trajectories^66,67^to compute residue-based stability profiles. The fluctuations of the mean distance between a given residue and all other residues in the ensemble were converted into distance fluctuation stability indexes that measure the energy cost of the residue deformation during simulations. The high values of distance fluctuation indexes are associated with stable residues that display small fluctuations in their distances to all other residues, while small values of this stability parameter would point to more flexible sites that experience large deviations of their inter-residue distances. The distance fluctuation stability index for each residue is calculated by averaging the distances between the residues over the simulation trajectory using the following expression:

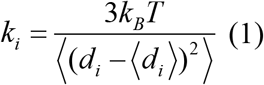

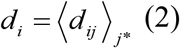

*d*_*ij*_ is the instantaneous distance between residue *i* and residue *j, k*_*B*_ is the Boltzmann constant, *T* =300K. ⟨ ⟩ denotes an average taken over the MD simulation trajectory and 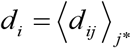 is the average distance from residue *i* to all other atoms *j* in the protein (the sum over *j*_*_ implies the exclusion of the atoms that belong to the residue *i*). The interactions between the *C*_*α*_ atom of residue *i* and the *C*_*α*_ atom of the neighboring residues *i* −1 and *i* +1 are excluded in the calculation since the corresponding distances are nearly constant. The inverse of these fluctuations yields an effective force constant *k*_*i*_ that describes the ease of moving an atom with respect to the protein structure.

### Mutational Scanning and Sensitivity Analysis

We conducted mutational scanning analysis of the binding epitope residues for the SARS-CoV-2 S protein complexes. Each binding epitope residue was systematically mutated using all 19 substitutions and corresponding protein stability changes were computed. BeAtMuSiC approach^68-70^ was employed that is based on statistical potentials describing the pairwise inter-residue distances, backbone torsion angles and solvent accessibilities, and considers the effect of the mutation on the strength of the interactions at the interface and on the overall stability of the complex. The binding free energy of protein-protein complex can be expressed as the difference in the folding free energy of the complex and folding free energies of the two protein binding partners:

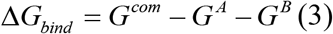

The change of the binding energy due to a mutation was calculated then as the following:

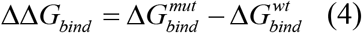

We leveraged rapid calculations based on statistical potentials to compute the ensemble-averaged binding free energy changes using equilibrium samples from MD trajectories. The binding free energy changes were computed by averaging the results over 1,000 equilibrium samples for each of the studied systems.

### Dynamic Network Analysis

A graph-based representation of protein structures^71,72^ is used to represent residues as network nodes and the inter-residue edges to describe non-covalent residue interactions. The network edges that define residue connectivity are based on non-covalent interactions between residue side-chains that define the interaction strength *I*_*ij*_ according to the following expression used in the original studies.^71,72^

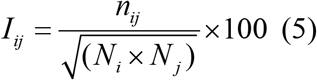

where *n*_*ij*_is number of distinct atom pairs between the side chains of amino acid residues *i* and *j* that lie within a distance of 4.5 Å. *N*_*i*_ and *N* _*j*_ are the normalization factors for residues *i* and *j*. We constructed the residue interaction networks using both dynamic correlations^73^ and coevolutionary residue couplings^74^ that yield robust network signatures of long-range couplings and communications. The details of this model were described in our previous studies.^75-77^

In brief, the edges in the residue interaction network are then weighted based on dynamic residue correlations and coevolutionary couplings measured by the mutual information scores. The edge lengths in the network are obtained using the generalized correlation coefficients ***R***_*MI*_ (***X***_***i***_, ***X***_***j***_) associated with the dynamic correlation and mutual information shared by each pair of residues. The length (i.e. weight) *w*_*ij*_ = − log[***R***_*MI*_(***X***_***i***_, ***X***_***j***_)] of the edge that connects nodes *i* and *j* is defined as the element of a matrix measuring the generalized correlation coefficient ***R***_*MI*_(***X***_***i***_, ***X***_***j***_) as between residue fluctuations in structural and coevolutionary dimensions. Network edges were weighted for residue pairs with ***R***_*MI*_(***X***_***i***_, ***X***_***j***_) > 0.5 in at least one independent simulation as was described in our initial study. The matrix of communication distances is obtained using generalized correlation between composite variables describing both dynamic positions of residues and coevolutionary mutual information between residues. As a result, the weighted graph model defines a residue interaction network that favors a global flow of information through edges between residues associated with dynamics correlations and coevolutionary dependencies. To characterize allosteric couplings of the protein residues and account for cumulative effect of dynamic and coevolutionary correlations, we employed the generalized correlation coefficient first proposed by Lange and Grubmüller.^78^ The g_correlation tool in the Gromacs 3.3 package was used that allows computation of both linear or non-linear generalized correlation coefficients.^79^ The protocol was detailed in our earlier study^74^ showing that the generalized correlation coefficient based on dynamic and coevolutionary can accurately determine the cross-correlation between protein residues. This approach has also been utilized in a similar context of allosteric modeling by other groups^80-83^ where the generalized correlation GC matrix measured dynamic interdependencies between on fluctuations of spatially separated residues.

The RING program^84,85^ was also employed for the initial generation of residue interaction networks. The ensemble of shortest paths is determined from matrix of communication distances by the Floyd-Warshall algorithm.^86^ Network graph calculations were performed using the python package NetworkX.^87^

The Girvan-Newman algorithm^88,89^ is used to identify local communities. In this approach, edge centrality (also termed as edge betweenness) is defined as the ratio of all the shortest paths passing through a particular edge to the total number of shortest paths in the network. The method employs an iterative elimination of edges with the highest number of the shortest paths that go through them. By eliminating edges the network breaks down into smaller communities. The algorithm starts with one vertex, calculates edge weights for paths going through that vertex, and then repeats it for every vertex in the graph and sums the weights for every edge. However, in complex and dynamic protein structure networks it is often that number of edges could have the same highest edge betweenness. An improvement of Girvan-Newman method was implemented and the algorithmic details of this modified scheme were given in our recent studies.^90,91^ Briefly, in this modification of Girvan-Newman method, instead of a single highest edge betweenness removal, all highest betweenness edges are removed at each step of the protocol. This modification makes community structure determination invariant to the labeling of the nodes in the graph and leads to a more stable solution. The modified algorithm proceeds through the following steps: a) Calculate edge betweenness for every edge in the graph; b) Remove all edges with highest edge betweenness within a given threshold; c) Recalculate edge betweenness for remaining edges; d) Repeat steps b-d until graph is empty.

The betweenness of residue *i* is defined as the sum of the fraction of shortest paths between all pairs of residues that pass through residue *i*:

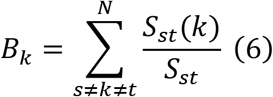

where *S*_*st*_ denotes the number of shortest geodesics paths connecting s and *t*, and *S*_*st*_(*k*) is the number of shortest paths between residues *s* and *t* passing through the node *k*.

The following Z-score is then calculated

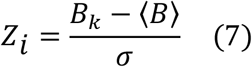

Through mutation-based perturbations of protein residues we compute dynamic couplings of residues and changes in the average short path length (ASPL) averaged over all possible modifications in a given position. The change of ASPL upon mutational changes of each node is inspired and reminiscent to the calculation proposed to evaluate residue centralities by systematically removing nodes from the network.

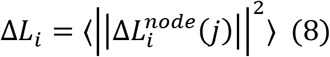

where *i* is a given site, *j* is a mutation and⟨⋯⟩denotes averaging over mutations. 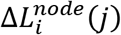 describes the change of ASPL upon mutation *j* in a residue node *i*. Δ*L*_*i*_ is the average change of ASPL triggered by complete mutational profiling of this position.

Z-score is then calculated for each node as follows:

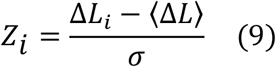

⟨Δ*L*⟩ is the change of the ASPL under mutational scanning averaged over all protein residues in the S-RBD and σ is the corresponding standard deviation. The ensemble-averaged Z –scores ASPL changes are computed from network analysis of the conformational ensembles using 1,000 snapshots of the simulation trajectory for the native protein system.

## SUPPORTING INFORMATION

Supporting information contains three additional figures showing ensemble-averaged conformational mobility profiles (Figure S1), SASA profiles (Figure S2) and structural maps of allosteric communication hotspot residues for the SARS-CoV-2 S timer complexes (Figure S3).

## AUTHOR INFORMATION

The authors declare no competing financial interest.

## ACKNOWLEDGMENT

This work was supported by the institutional funding from Chapman University. The author also acknowledges support by the Kay Family Foundation Grant A20-0032.

## Notes

### Competing Interest Statement

The authors have declared no competing interest.

