## Supporting Figures S1-S3 for "Deep Mutational Scanning of Dynamic Interaction Networks in the SARS-CoV-2 Spike Protein Complexes: Allosteric Hotspots Control Functional Mimicry and Resilience to Mutational Escape"

*Gennady M. Verkhivker,*[1,2] *

[1]Keck Center for Science and Engineering, Department of Computational and Data Sciences, Schmid College of Science and Technology, Chapman University, One University Drive, Orange, CA 92866, USA

[2] Depatment of Biomedical and Pharmaceutical Sciences, Chapman University School of Pharmacy, Irvine, CA 92618, USA

***Corresponding Author**

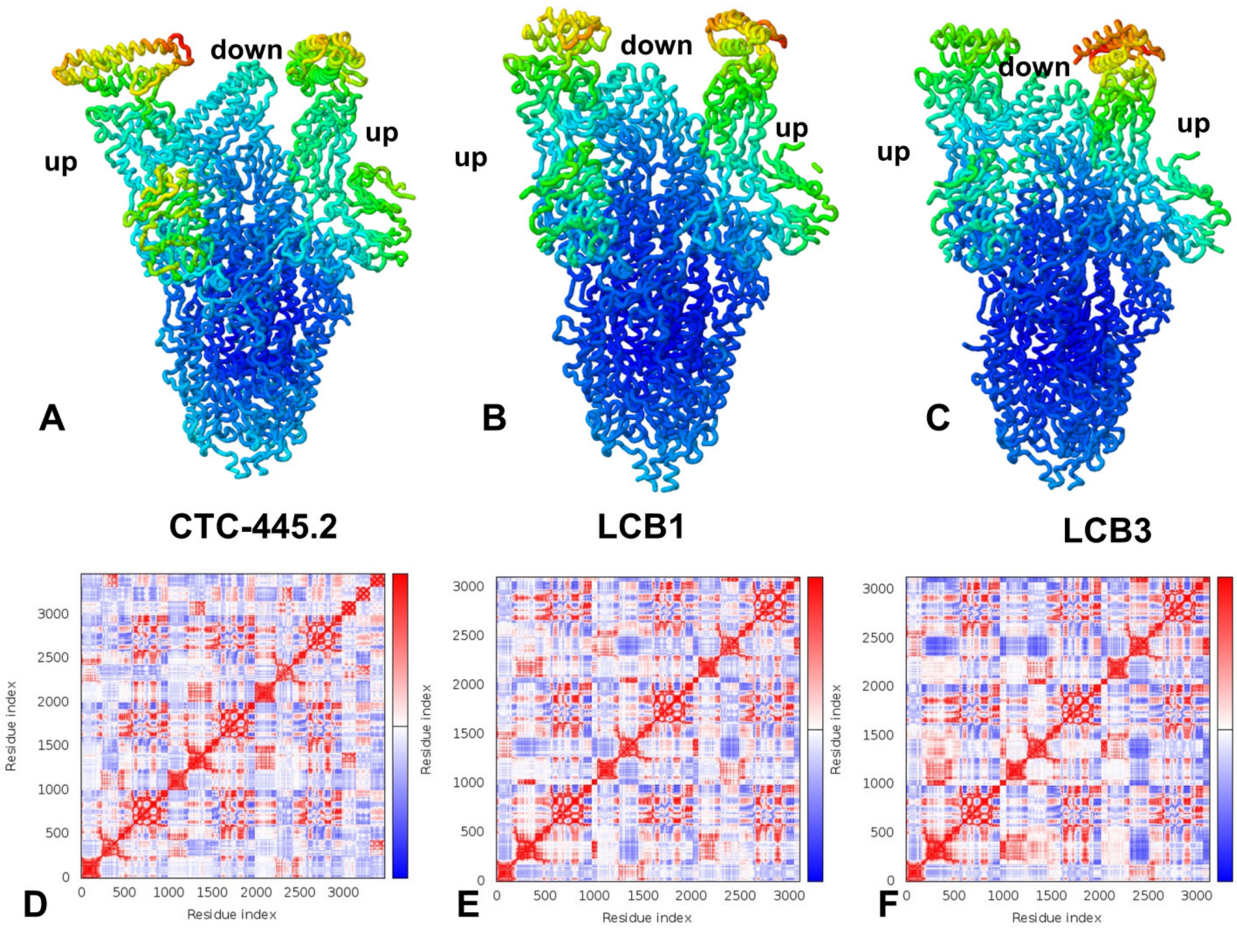


**Figure S1.** Conformational mobility profiles for the SARS-CoV-2 S timer complexes with CTC-445.2 decoy, pdb id 7KL9 (A), complex with LCB1 miniprotein, pdb id 7JZL (B), and complex with LCB3 miniprotein, pdb id 7JZN (C). The structures are in ribbons with the rigidity-to-flexibility scale colored from blue to red**.** (D-F) The covariance matrix indicates coupling between pairs of residues. Cross-correlations of residue-based fluctuations vary between +1 (correlated motion; fluctuation vectors in the same direction, colored in dark red) and -1 (anti-correlated motions; fluctuation vectors in the same direction, colored in dark blue). The values > 0.5 are colored in dark red and the lower bound in the color bar indicates the value of the most anti-correlated pairs.

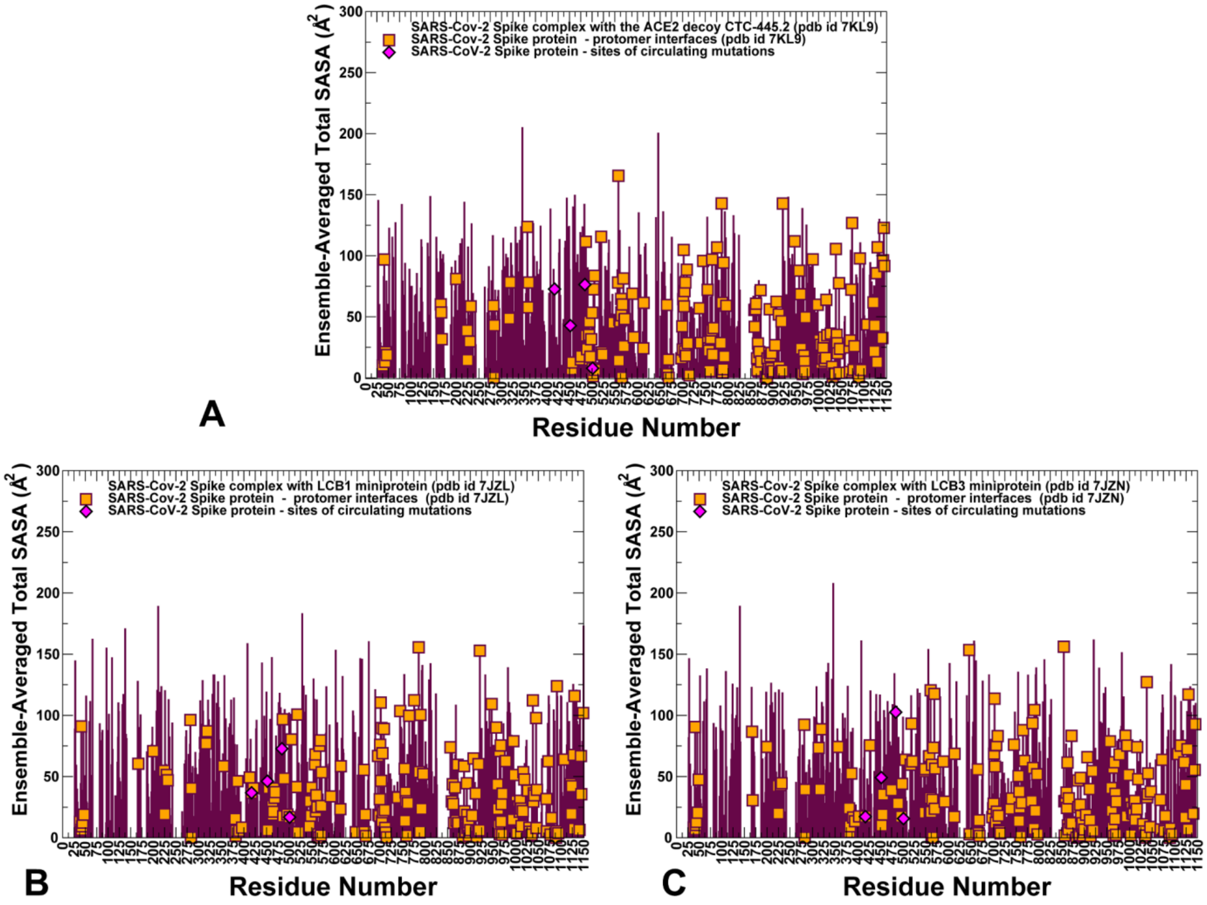


**Figure S2.** The ensemble-average average total solvent accessible surface area (SASA) for the SARS-CoV-2 S timer complexes with CTC-445.2 decoy, pdb id 7KL9 (A), complex with LCB1 miniprotein, pdb id 7JZL (B), and complex with LCB3 miniprotein, pdb id 7JZN (C). The SASA profiles are shown in maroon lines. The protomer residues involved in both protomer-protomer contacts and intermolecular interactions with bound miniproteins are shown by orange-filled squares. A The positions of functional RBD sites subjected to circulating mutational variants K417, L452, E484, and N501 are highlighted in magenta-colored filled diamonds.

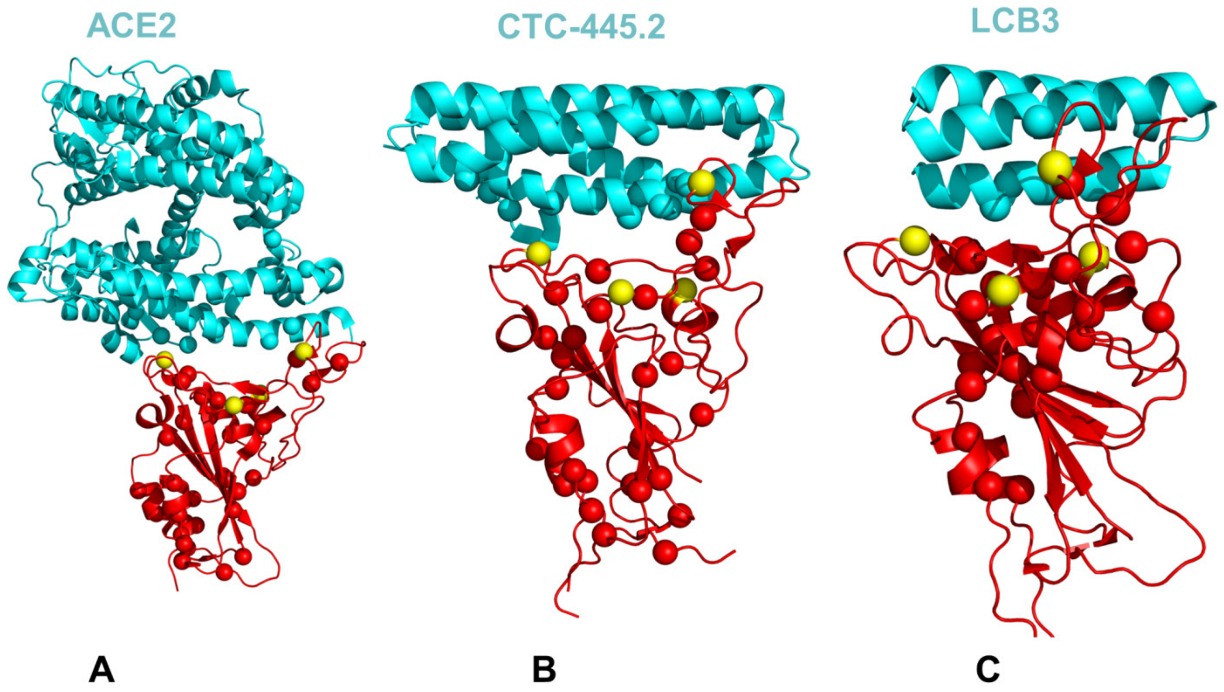


**Figure S3.** Structural maps of allosteric communication hotspot residues for the SARS-CoV-2 S timer complexes with CTC-445.2 decoy, pdb id 7KL9 (A), complex with LCB1 miniprotein, pdb id 7JZL (B), and complex with LCB3 miniprotein, pdb id 7JZN (C). The hotspot residues highlight peaks of the distribution depicting the ASPL changes induced by mutations in respective protein residues. By identifying residues in which mutations cause a significant change of the ASPL parameter, this network-based mutational profiling identifies allosteric hotspot positions that are important for signal transmission in the SARS-CoV-2 complexes. The S-RBD is shown in red ribbons. The allosterically important residues are shown in red-colored spheres. The functional sites K417, L452, E484, and N501 are highlighted in yellow spheres. The binding proteins ACE2, CTC-445.2 and LCB3 are shown in cyan colored ribbons and annotated.
